# Effects of transcranial focused ultrasound stimulation to human lateral geniculate nucleus on visual perception and steady-state visual evoked potentials

**DOI:** 10.64898/2026.07.30.741804

**Authors:** Martin TW Scott, Patti N Limon, Morteza Mohammadjavadi, Benjamin R. Kop, Nai-Feng Chen, Eva A Feredoes, Vladimir Vildavski, Gerald R Popelka, Anthony M. Norcia, Kim Butts Pauly, Ryan T Ash

## Abstract

Transcranial ultrasound stimulation (TUS) is an emerging tool to non-invasively modulate neural activity in deep brain areas. A key need in accelerating TUS into cognitive neuroscience and neuropsychiatry is to better understand how different sonication parameters relate to neuromodulatory effects. Here we assess the role of pulse repetition frequency (PRF), a key TUS parameter thought to determine the relative contribution of molecular displacement and acoustic radiation force effects on neural tissue using the human subcortical visual pathway as a testbed. We combined frequency-tagged steady-state visual evoked potential (SSVEP) measures of contrast-response with contrast increment detection psychophysics as neural and behavioral readouts of visual pathway function. We used structural MRIs and acoustic simulations to target the lateral geniculate nucleus (LGN). Concurrent with visual stimulus presentation, the left LGN or a more superficial control site were stimulated with a neuronavigated depth-steerable 4-element TUS transducer at a range of PRFs with 68 W/cm^2^ free-water I_SPPA_, and a 10% duty cycle. An effective white-noise auditory mask blinded participants to stimulation conditions. Recordings from 25 neurotypical participants failed to detect any impact of TUS on SSVEP response amplitude, SSVEP response latency, or perceptual behavior. Analysis of simulations generated from the measured transducer positions grant reasonably high confidence that the LGN was within the TUS focus in most participants, with no correlation between targeting accuracy and changes in activity during TUS. Our results provide a cautionary note about the effect size of neuronavigated TUS for online causal manipulations in cognitive and clinical neuroscience.

## Introduction

Transcranial ultrasound stimulation (TUS) is an emerging technology that enables targeted noninvasive neuromodulation of deep brain structures in humans (Osada and Konishi 2024). TUS involves the focusing of ultrasound waves originating from ultrasonic transducers placed on the scalp to modulate nervous system function. Amongst non-invasive neuromodulation techniques, TUS has the unique advantage of a highly focal modulation volume (as small as ∼3 mm^3^) that can be steered to different target anatomical structures (Martin et al. 2025). Recent research has demonstrated that TUS can produce reversible changes in neural circuit activity and behavior at safe ultrasound intensities in preclinical models (Menz et al. 2013; Dallapiazza et al. 2018; Folloni et al. 2019; Fouragnan et al. 2019; Munoz et al. 2022; Murphy et al. 2022, 2024; Niu, Yu, and He 2022; Di Ianni et al. 2023; Lu et al. 2024) and in humans (Legon et al., 2018; Butler et al., 2022; Yaakub et al., 2023, 2024; Bancel et al., 2024; Bao et al., 2024; Chou et al., 2024; Strohman et al., 2024; Zadeh et al., 2024; Barksdale et al., 2025; Caulfield et al., 2025; Farboud et al., 2025). The underlying neuromodulatory mechanism is presumed to occur via the opening of endogenously mechanosensitive ion channels that are highly sensitive to ultrasound, with the effect of increasing or decreasing neuronal excitability (Yoo et al., 2022). However, there also is evidence that the mechanical effects of TUS primarily modulate non-neuronal cells, including glia and vascular cells (Newman et al. 2024), and that the effects emerge gradually over the course of hours. Some have hypothesized that the immediate online effects of TUS can sometimes be primarily mediated through a thermal mechanism (Darrow et al. 2019) and/or unintended auditory co-stimulation (Kop et al. 2024).

The optimal online stimulation parameters to suppress or enhance activity have not yet been established in humans (Nandi et al. 2024), and currently TUS does not have a high-throughput method akin to a primary motor response induced by transcranial magnetic stimulation (TMS) to evaluate the effect of different stimulation parameters (Säisänen et al. 2008). Targeting TUS to early sensory areas has shown initial promise to achieve this capability: TUS to the visual thalamus (lateral geniculate nucleus, LGN) was the first demonstration of low-intensity focused ultrasound neuromodulation, shown by the Fry brothers in cat (Fry, Ades, and Fry 1958). Later studies showed reversible changes in visual perceptual behavior in macaques with LGN TUS (Webb et al. 2022, 2023). Our lab also showed TUS neuromodulation effects in LGN of sheep (Mohammadjavadi et al. 2022). Recent studies have shown encouraging demonstrations of TUS effects in nonvisual thalamic areas (Caulfield et al. 2025; Legon et al. 2018), and a recent study suggests that TUS to the human LGN can modulate visual cortical blood oxygen level-dependent (BOLD) functional magnetic resonance imaging (fMRI) responses (Martin et al. 2025).

Expanding upon this work, we developed a novel paradigm to evaluate TUS parameters in human using the LGN as a testbed and use it to determine its impact on both electrophysiology and perceptual behavior. The LGN has many supportive features as a testbed for TUS: 1) It is small (<0.5 cm diameter), on the order of the TUS focus, so most of the nucleus can be modulated; 2) it is deep in the brain, allowing an evaluation of efficacy at depths that are clinically relevant (>6 cm); 3) it relays visual information from the retina to the primary visual cortex, so TUS-induced changes in LGN excitability can be read out in the occipital lobe with electroencephalography (EEG); 4) it has lateralized anatomy, in which the right visual field is represented in the left LGN and visual cortex, and vice versa, allowing rigorous internal control recordings in the ipsilateral non-stimulated hemifield. This half-visual field representation of each LGN can be leveraged by the steady-state visually evoked potential (SSVEP) paradigm, where stimuli in both hemifields can be simultaneously flickered at different frequencies, with the hemifield/LGN specific neural responses later separated with spectrum analysis (Norcia et al. 2015).

Our experimental design had four innovations that allow us to detect veridical TUS neuromodulation effects with less risk for auditory and expectancy confounds that could otherwise have a significant impact (Guo et al. 2018; Fong et al. 2025; Sato, Shapiro, and Tsao 2018; Kop et al. 2024): 1) The frequency-tagged visual stimuli were presented at temporal frequencies that did not overlap or share harmonics with the TUS pulse repetition frequencies or harmonics used. By quantifying neural responses at the frequencies of the visual stimulus, we avoided contribution from any TUS-evoked auditory or somatosensory responses (Kop et al. 2025) or potential TUS electrical artifacts; 2) responses in the ipsilateral non-sonicated hemifield provided a rigorous internal control measure that permits detection of non-specific TUS effects; 3) the depth of the TUS focus was varied on each trial to target the LGN (the transducer’s maximum depth; 70 mm) or a control site in the more superficial auditory cortex and subcortical white matter (the transducer’s minimum TUS depth; 30 mm), which matched the sensorial co-stimulation of the active condition but avoided the main visual pathways; 4) a highly-effective white noise mask rendered the TUS auditory co-stimulation inaudible in the majority of participants, ensuring both participant and experimenter blinding between the conditions. Therefore, if TUS is effective and focal, it should have a lateralized effect that is more prominent in the sonicated hemifield, and TUS to the LGN should have more pronounced effects than control-site TUS.

We used our approach to test the role of pulse repetition frequency (PRF) in TUS neuromodulation effects. PRF is a key parameter because, for the low duty cycles typically used in TUS, it controls the duration of individual pulses. Long pulses and short pulses will elicit different proportions of acoustic radiation force and molecular displacements. There is preliminary evidence that long pulses may be more excitatory, while short pulses may be more inhibitory (Nandi et al. 2024). In contrast to prior LGN TUS studies (Fry, Ades, and Fry 1958; Mohammadjavadi et al. 2022; Martin et al. 2025) our experiment failed to detect significant effects of ultrasound on either electrophysiology or perceptual behavior.

## Methods

### Participants

Healthy participants (n=25, M=14, F=13) ages 18-55 (mean=33, range=22-58) were recruited from the research community and Stanford University paid participant pool. Inclusion/exclusion criteria included corrected-to-normal vision, no MRI contraindications, no severe neurological disorder or epilepsy, no age-related hearing loss, and no history of intracranial mass. Participants were consented following Stanford University Institutional Review Board guidelines. Participants were paid $30 per hour. Twenty-five subjects were retained with 2 rejected due to poor SSVEP signal quality.

### General approach

Participants were invited for two separate experimental sessions. **Figure 1** provides an overview of the contents of each session for a single participant. First, participants attended a neuroimaging session where anatomical MRI scans were collected for LGN segmentation and TUS planning. TUS simulations were conducted after session 1, in preparation for session 2. Session 2 had several stages, as shown in **Figure 1**. Initially, participants took part in a brief high-density EEG recording while viewing the SSVEP- eliciting stimulus. The high-density recording was immediately analyzed, and the SSVEP topography was used to guide the later placement of three EEG channels at scalp sites identified as having high signal levels for SSVEP. Subsequently, the participant was positioned in the neuronavigation system. The participant’s hair was prepared with ultrasound gel, the pre-specified EEG electrodes were placed, and the TUS transducer was firmly mounted at the pre-planned location. For all following stages, both SSVEP and behavioral data were collected, and the auditory mask was added. These stages were a pre- TUS baseline stage, followed by the ‘core conditions’ stage (where TUS was switched on with different depth and PRF settings), and ending with the post-TUS stage. All data were normalized to the data collected in the pre-TUS baseline stage (details to follow).

**Figure 1:**
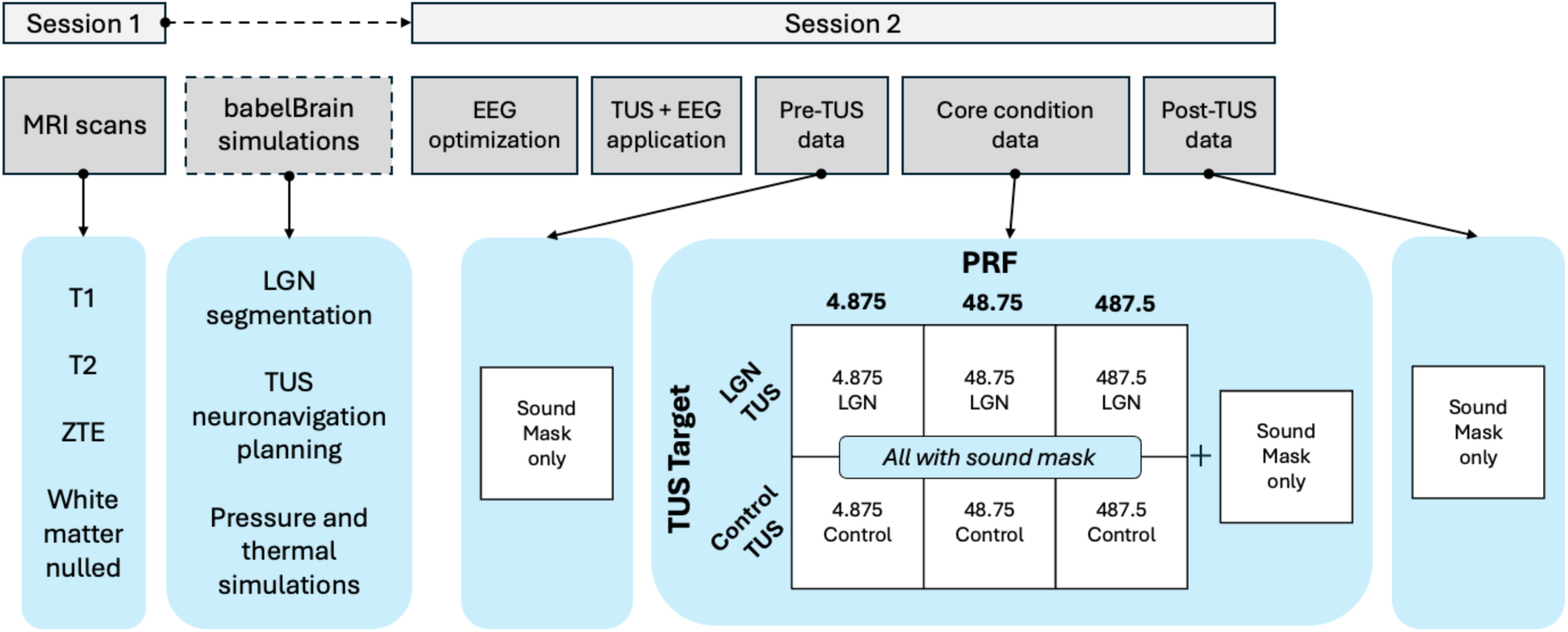
Schematic overview of the study progression. The middle darker-grey row illustrates the experimental stages. The blue shaded boxes include detailed information for each stage.

### MRI Neuroimaging

MR images were acquired on a 3T scanner (DV750, GE Healthcare, Milwaukee, WI) with spin echo (SE) sequences. An 8-channel head coil (MRI Devices) was used. Four MRI protocols were acquired with the following scan parameters, with a 240×240 mm field-of-view. **<u>T1</u>:** TR/TE: 5.6ms/2.3ms, BW: 62.5Hz, FOV: 24cm, slice thickness: 0.8mm, matrix size: (freq:300, phase:256). **<u>T2</u>:** TR/TE: 2509ms / 120ms, BW: 62.5Hz, FOV: 24cm, slice thickness: 0.8mm, matrix size: (300,256). **<u>ZTE</u>:** TR/TE: 37ms / 0.016ms, BW: 62.5Hz, FOV: 24cm, slice thickness: 0.8mm, matrix size: (300, 300). **<u>WMN</u>:** TR/TE: (9.13/8.6ms) / 3.9ms, BW: 25Hz, FOV: 24cm, slice thickness: 0.8mm, matrix size: (300,180).

### Trajectory planning and simulations

A biophysical head model was generated with the SimNIBS charm function (Puonti et al. 2020). The LGN ROI was determined with hipTHOMAS software (Vidal et al. 2024; Su et al. 2019) applied to the WMN scan. The SimNIBS head model and LGN ROI were imported into Brainsight software (Rogue Research Inc., Montreal). The TUS trajectory was placed at the center of the left LGN ROI, oriented to minimize the distance from the scalp to the target, attain a near-perpendicular angle of incidence relative to the scalp, and to maintain >3 cm distance (the transducer radius) from the participant’s external ear. For most participants, the ideal transducer position to minimize trajectory angle would collide with the external ear, so the transducer had to be translated slightly, such that for 22 participants the final transducer position was immediately superior to the external ear, and with 3 participants the transducer was anterior and slightly superior to the external ear. The angle of the transducer relative to the head was 5-10°. Trajectories were simulated in open-source focused ultrasound simulation software (BabelBrain (Pichardo 2023)), at 6 points per wavelength (PPW), using each individual’s SimNIBS head model, ZTE MR skull image, and personalized LGN trajectory. Z-axis steering of 70 mm was used with a scalp offset of 7 mm (to account for the gel pad), except in two participants with smaller heads, for whom 62 and 65 mm depth was used. The ‘mechanical adjustment’ function of the BabelBrain software was used to slightly reposition the transducer to account for the acoustic lensing properties of the skull. Thermal simulations were performed in BabelBrain, which showed a maximum temperature rise of 0.5° C at the target, and CEM43 thermal dose < 0.25, following International Consortium for Transcranial Ultrasonic Stimulation Safety and Standards guidelines (Aubry et al. 2025). In the post-hoc simulations to estimate dose at the target, the most common transducer position was identified from the sampled transducer position during TUS that was recorded online in BrainSight. The simulation was performed centered at 70 mm deep to the transducer aperture (or shallower in the two participants with smaller heads). The distance from the scalp was measured for each participant in Brainsight and entered into the simulation software.

### EEG preparation and recording

EEG was collected using two different montages: a 128-channel high density montage and a 3- channel sparse montage. The high-density montage was used to find the EEG channels with the highest visually evoked signal for each checkerboard reversal frequency. This high-density recording was collected and analyzed 20 minutes prior to each participant’s experimental ultrasound session. The sparse montage was used in lieu of a custom high-density montage to leave room for the TUS transducer to be mounted to the participant’s head. The placement of the sparse montage was customized for each participant to match the location of peak amplitude channels found in the high-density recording. The location of the reference channel (Cz) and ground was kept consistent across both montages. The high-density recording used a saline-based net system (EGI HydroCel GSN 130) while the low-density used gold-cup electrodes and a 4 channel amplifier (A-M Systems model 1700). For the low-density recording, electrode sites were prepared with abrasive gel (Nuprep), and electrodes were coupled to the scalp with conductive paste (Ten20). The locations of the low-density electrodes were digitized in BrainSight following neuronavigation. EEG quality was confirmed by observation of eyes-closed alpha oscillations on the in the live-sampled data. EEG was sampled by in-house software (XDiva) at 420 Hz (7 data samples per video frame).

### Neuronavigation and TUS preparation

A frameless neuronavigation head tracker (Rogue Research, inc.) was adhered to the participant’s forehead with double-sided body tape. The participant’s head fiducials (left and right tragus or preauricular point, left and right outer canthus, and nasion) were marked in Brainsight (Rogue Research, Inc.). During the validation step of head registration, additional corrective points were placed as needed such that the pointer-to-scalp error was <2 mm on the left, right, top, and front of the head. Participants sat in a Brainsight TMS chair (Rogue Research, Inc.) with their head supported in three places to minimize movement: a rest for the chin, head support on the right, and seat cushion on the back of the head. The transducer had an attached neuronavigation tracker that was calibrated in Brainsight at the beginning of each experiment session. The transducer was initially placed without gel at the simulation- determined trajectory, and the scalp at the edges of the transducer were marked with a wax pencil. The transducer was removed, and the hair was prepared following established guidelines (Murphy et al., 2025). In brief, the hair was infiltrated with ultrasound gel and parted in layers starting from the outer edge of the transducer location and moving inward. For each layer of hair, gel was infused into the hair with finger pressure. Repeated application of gel allowed bubbles to be palpated and extruded. The goal of the preparation was to minimize the presence of gas bubbles that could interfere with ultrasound propagation and to align the hair in an orderly, layered fashion to further support good ultrasound propagation. A generous amount of gel was applied to the transducer face ensuring no deposited bubbles, and a custom molded transparent gel pad with a precut angle to accommodate the desired trajectory (5 or 10° from perpendicular incident angle), 5 mm at its thinnest point, was fitted into the transducer face. This gel pad had a convexity that near-perfectly matched the concavity of the transducer and once applied to the gelled transducer it was carefully massaged to remove any remaining bubbles. The transducer and gel pad stack were then affixed to the left side of the participant’s head at the planned neuronavigation trajectory and immobilized with an articulated arm (Rogue Research, Inc.). Light pressure was applied to maintain contact between the gel pad and scalp. The transducer was positioned to match the angle and position determined by simulations to target the LGN. The transducer position was continually monitored throughout the experimental session to ensure the TUS focus remained on the target. If the focus moved >1.5 mm from the target based on the live neuronavigation, the experiment was paused and the transducer was repositioned, and the TUS transducer position was logged online in Brainsight for later post-hoc simulations.

### TUS parameters

We used the NeuroFUS Pro CTX-500 system (supplier/support: BrainBox Ltd., Cardiff, UK; manufacturer: Sonic Concepts Inc., WA, USA) with a 500 kHz acoustic center frequency, 64mm spherical radius, 64mm aperture diameter, 4-element annular transducer. The transducer had a depth range of 30 to 70 mm. We measured the focus by hydrophone in a scan tank and confirmed that the transducer had a focal spot of 0.5 x 0.5 x 2 cm full width at half maximum (FWHM), extended as expected in the axial dimension (**Supp. Fig. 1**). Transducer output was periodically measured by hydrophone throughout the course of experimentation and was found to be stable throughout. Ultrasound was delivered in 12 second trains overlapping in time with the SSVEP eliciting stimuli. TUS was triggered by TTL pulse to start simultaneously with visual stimulus onset (NeuroFUS trigger mode 4, microsecond trigger precision). Free-water ultrasound spatial peak pulse averaged intensity (I_SPPA_) was set to 68 W/cm^2^ (free-water acoustic pressure: 1.4 MPa) for all participants. Per BabelBrain simulations, the estimated in situ I_SPPA_ was 16.2±3.6 W/cm^2^ (range 8.5-20.5), equivalent to a peak negative acoustic pressure of 0.69±0.33 MPa (range 0.5-0.78). Duty cycle (DC) was maintained at 10%. PRF was randomized on a trial-by-trial basis among 4.875 Hz, 48.75 Hz, and 487.5 Hz. These PRFs were chosen such that there would be an integer number of pulse cycles during each 11s trial. This ensures that any neural activity as a direct consequence of TUS stimulation is centered into a Fourier spectrum bin, thus increasing the probability of detecting this signal if it is present. Additionally, these PRFs were chosen to minimize the Fourier bin overlap between direct TUS-evoked activity harmonics and with the harmonics we expect visual stimulus. TUS pulses were linear-ramped for the first and last 5% of the pulse duration to decrease audibility for the two lower PRFs. Two different depths were used for the TUS focus, 70mm (LGN-TUS) and 30mm (Control-TUS), as shown in **Figure 2A** (apart from two aforementioned exceptions for participants with a particularly shallow LGN).

**Figure 2.**
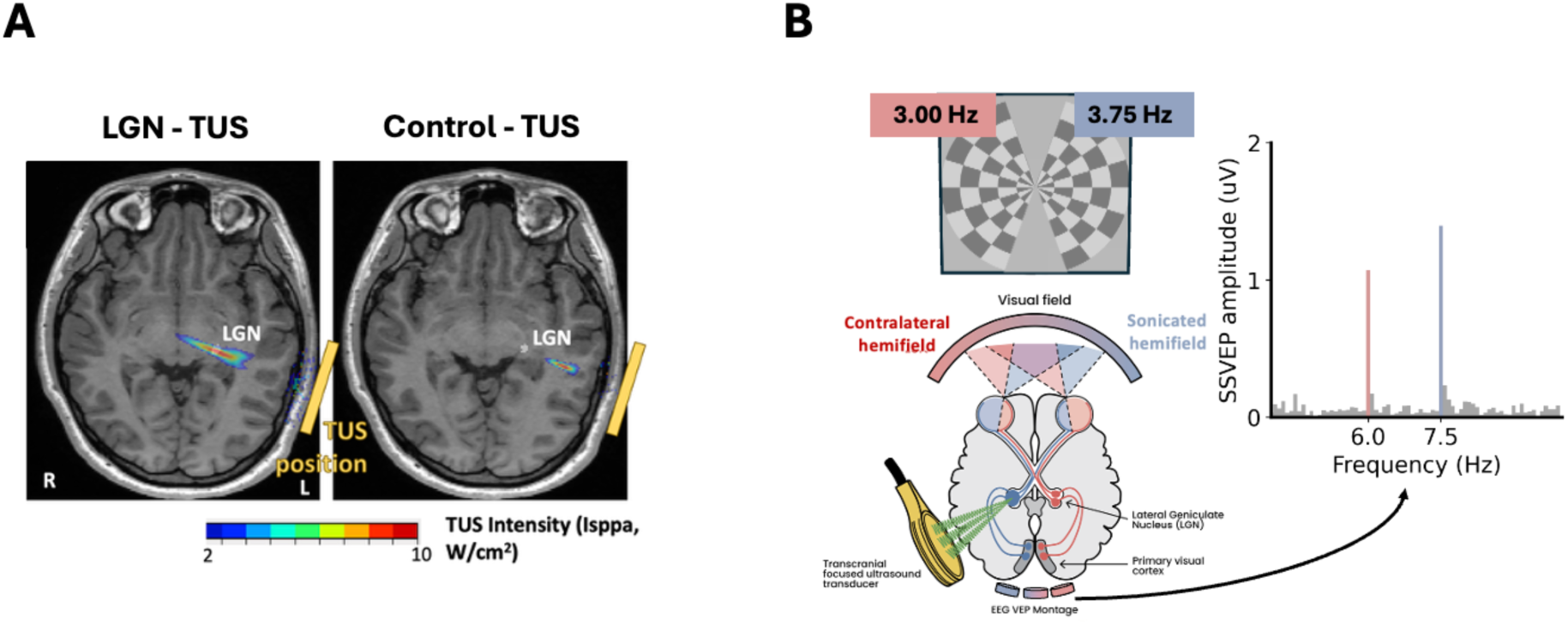
Illustration of SSVEP visual stimulation paradigm and TUS depth-based control. **A**. Horizontal sections of MRI through the LGN of an example participant, showing heatmap depictions of the simulated TUS intensity in the LGN TUS condition, with depth steered to 70 mm to target the LGN, and the Control TUS condition, with depth steered to 30 mm, targeting subcortical white matter generally near auditory cortex. Images shown in radiological convention, with left hemisphere on the right. **B**. Schematic showing the hemifield representation of visual information by the visual system, targeting of TUS to left LGN, and sparse occipital EEG montage. Inset on right shows an example participant’s EEG Fourier spectrum showing SSVEP amplitude at the 2F frequency tags of each hemifield visual stimulus.

### Visual stimuli

Following Norcia et al., (2015) and Ash, Nix, and Norcia., (2024) frequency-tagged checkerboard wedge stimuli, generated with in-house xDIVA software, were presented on a 60-Hz refresh-rate monitor (Dell G2524H) positioned 80 cm from the participant’s eye level. Each trial consisted of two contrast-reversing checkerboard wedges presented in the left and right hemifields at two different full-cycle frequencies 3 Hz in the left hemifield (F1), and 3.75 Hz in the right hemifield (F2) (**Fig. 2B**). Sign-reversing checkerboards generate neural responses at the reversal frequency (twice the full-cycle frequency), such that neural responses are expected at 6 Hz (2F2) and 7.5 Hz (2F2) respectively (**Fig. 2B, inset,** example single-participant SSVEP Fourier spectrum shown), as well as the integer multiples of these frequences (4F2, 6F2, etc.). The stimuli subtended 0.5 to 10 degrees visual field angle. Each checkerboard wedge was divided into sixteen 0.628° segments along the vertical meridian relative to fixation, and 20 rings with a spatial frequency of 0.5°, creating a wedged checkerboard pattern. Stimuli flickered at 40% contrast and were presented in eight 1.33 second bins with 1.33 sec pre and post bins without TUS (13.333 seconds total, 10.667 sec TUS per trial). There was a 5 second intertrial interval. Participants were instructed to fixate at a central cross and minimize blinking during stimulus presentation. During the trial participants listened to a white-noise auditory mask. This auditory mask was played binaurally at a sound pressure level of 91dB(A) over a pair of Koss KSC75 ear-clip headphones which were calibrated using a sound level meter (Larson Davis SoundAdvisor 831c) with an ear simulator (Larson Davis AEC201-A). This mask level has been shown to drive TUS detection to near-chance level in the majority of participants (**Supp. Figure 2**).

### Behavioral task

During TUS-EEG, participants performed a psychophysical task requiring detection of a contrast increment of a 1° square on top of the frequency tagged SSVEP stimuli (**Fig. 3**), adapted from Lee (2023). The change could occur in the upper left, lower left, upper right, or lower right quadrant at 5° eccentricity. There were six behavioral probes presented per trial. The probes were always presented on the nearest rising edge of the square-wave checkerboard reversal. Participants were instructed to indicate (using a response keypad) whether they saw the probe in the upper or lower half of the whole checkerboard display. They were instructed not to base their response on whether the increment was in the left or right hemifield. The magnitude of the contrast increment was controlled by the QUEST+ adaptive search algorithm, with the lower asymptote fixed to chance (50%) and the upper asymptote fixed to 98% to allow for response errors (Watson 2017). Though participants were instructed to respond to every trial (even when they were uncertain), this task was not strictly 2-alternative forced choice as there was a finite 750ms behavioral response window after each probe. Sixty behavioral trials were recorded per condition.

**Figure 3:**
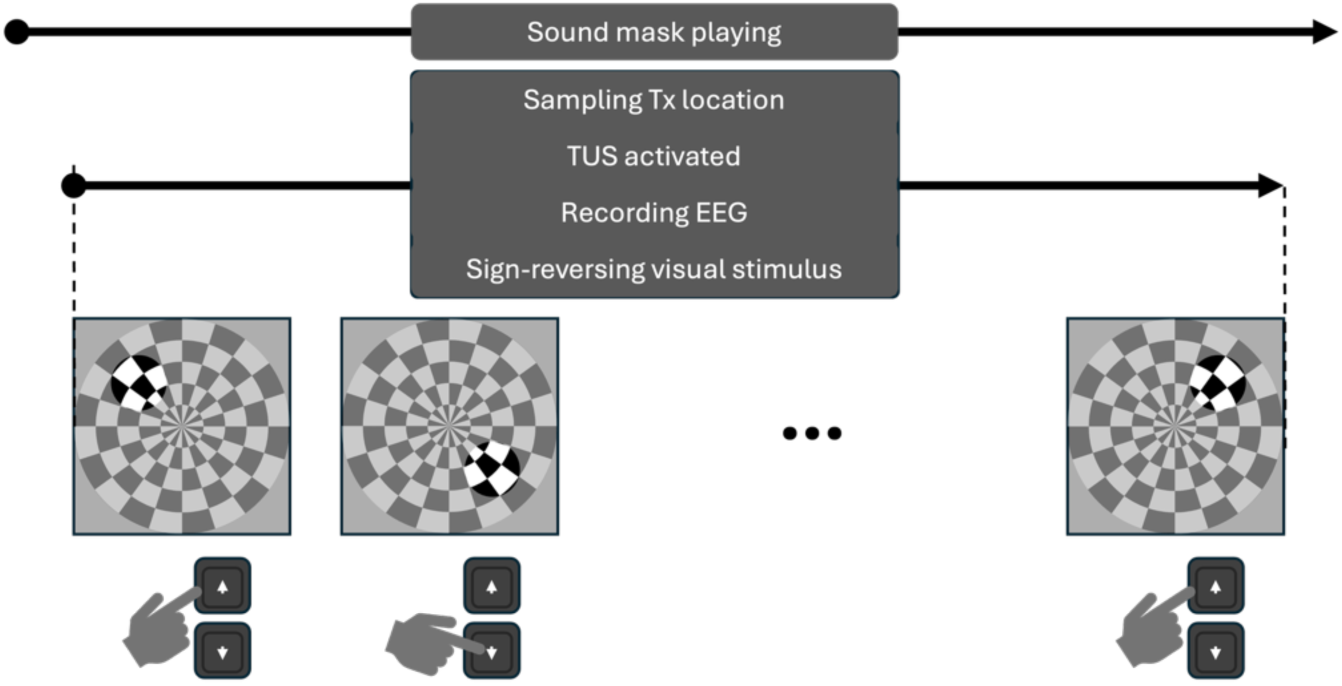
Illustration of the concurrent operations for a single trial in the ‘core conditions’ phase of this experiment.

### Experimental procedure

Prior to the start of the first task block, participants were advised that the TUS could generate sounds including beeps, buzzes, and tones, and they were also advised to notify the experimenter if they noticed any heating or discomfort at the TUS site. The experiment was a within-participants design with the following factors and levels: PRF (3 levels: 4.875 Hz, 48.75 Hz, 487.5 Hz), TUS target (2 levels: Control TUS, LGN TUS), and hemifield (2 levels: non-sonicated, sonicated, these data collected simultaneously). Participants first performed 16 no-TUS trials to establish a baseline. Then participants performed eight 16-trial TUS blocks with 2 randomly interleaved trials of each condition, with a total of 7 conditions (3 PRFs, 2 TUS targets + an interleaved No-TUS condition, as shown on Figure 1). Following the last TUS block, participants completed an additional 16 no-TUS trials. This final block of trials was used to investigate the presence of a hemifield-specific cumulative LGN-TUS neuromodulatory effect when compared to the pre-TUS baseline (i.e. an ‘offline effect’). No apparent visual percepts or discomfort at the TUS site were reported at any stage of the experiment.

### EEG Preprocessing

Due to the sparsity of the low-channel-count montage, we opted for a light-touch pre-processing approach. First, we applied 0.3 Hz – 50 Hz bandpass filter, then we removed the first 1.33 seconds of the EEG for every trial, for each condition (this period never contained TUS but did contain large visual stimulus onset transients). After this, for each participant, the amplitude and phase of the SSVEP were calculated at the 2^nd^, 4^th^, and 6^th^ harmonic by a recursive least squares (RLS) adaptive filter (Tang and Norcia 1995). The outputs of the RLS filter are similar to those of a Fourier transform, but RLS is more robust for short-duration samples. With our visual stimulus parameters, responses in most participants were restricted to the second harmonic of the reversal frequency (2F1 and 2F2), so our analysis was restricted to these components. For each condition, the complex-valued RLS coefficients were averaged across trials (coherent averaging), and these complex values were transformed to response amplitude and response phase. The SSVEP during LGN TUS was normalized by the SSVEP collected during the pre-TUS baseline within each hemifield. For amplitude measures, this was done by dividing a given condition by the amplitude at baseline, as divisive normalization is optimal for evoked potentials (Luck and Gaspelin 2017). For phase measures, the circular subtractive phase difference (relative phase) was calculated by subtracting the phase for a given condition from the phase for the baseline condition.

### Statistical analysis and psychometrics

Our primary hypothesis was that LGN TUS would have a different neuromodulatory effect than Control-TUS, and that the sonicated (right) hemifield would have more pronounced effects than the non-sonicated (left) hemifield. For the SSVEP data, we submitted the amplitude, phase, and complex coefficients (containing both amplitude and phase information) to within-subjects ANOVAs. Specifically, we used the robust resampling-based ANOVAs from the R-toolbox MANOVA.RM (Friedrich, Konietschke, and Pauly 2019), reporting the Wald-type statistic (WTS) rather than the F-test, with a permutation based p-value. For the behavioral data from each participant, and for each condition, a Weibull function was fit using the psychofit toolbox in Python (https://github.com/cortex-lab/psychofit). The slope and asymptote parameters of this fit were fixed to the same values as the QUEST+ search. The threshold search was initiated at the threshold obtained with QUEST+. The thresholds obtained by this fitting procedure were -log10 scaled to convert them to sensitivity (where higher is better), and the sensitivity values for all participants’ conditions were normalized (subtractive) to that of the pre-TUS baseline condition. These normalized values were submitted to a repeated-measures ANOVA. The SSVEP data were likewise normalized to the pre-TUS baseline condition, by dividing the amplitudes (in Microvolts) of all conditions by the amplitude of the pre-TUS baseline and subtracting the phase (in degrees) of all conditions from the phase of the phase of the pre-TUS baseline.

## Results

### Effect of LGN TUS on visual contrast increment detection sensitivity

We first tested the hypothesis that LGN TUS would disrupt or enhance visual perceptual sensitivity in the contralateral (sonicated) hemifield, and that this effect may be PRF-specific. The psychophysical sensitivity thresholds normalized to the pre-TUS baseline are shown in **Figure 4**. The hemifield (non-sonicated: top row, sonicated: bottom row) and PRF factor (4.8, 48.8, and 488 Hz from left to right) combinations are shown across plots, and the TUS targets (Control-TUS vs LGN-TUS) are shown within each plot, in addition to the mask-only condition (for reference). The 95% confidence intervals for perceptual performance overlap with each other across all experimental conditions, indicative of no statistically significant target-specific, hemifield-specific, or PRF-specific effect of TUS. There is a trend toward sensitivity being slightly below the unity line in all conditions, consistent with mild fatigue compared to the pre-TUS baseline to which all data were normalized. A repeated-measures ANOVA showed that there was no main effect of TUS target (WTS(1) = 2.16, p = 0.1522), no main effect of hemifield (WTS(1) = 0.01, p = 0.9142), and no main effect of PRF (WTS(2) = 0.08, p = 0.9602). Likewise, there were no significant interactions between TUS target and hemifield (WTS(1) = 0.09,p = 0.7606), target and PRF (WTS(2) = 0.80, p = 0.6869), or hemifield and PRF (WTS(2) = 0.82, p = 0.6789). The three-way interaction between target, hemifield, and PRF was also non-significant (WTS(2) = 0.10, p = 0.9558). Overall, this analysis provided no evidence that contrast increment detection thresholds differed as a function of target, hemifield, PRF, or any of their combinations.

**Figure 4:**
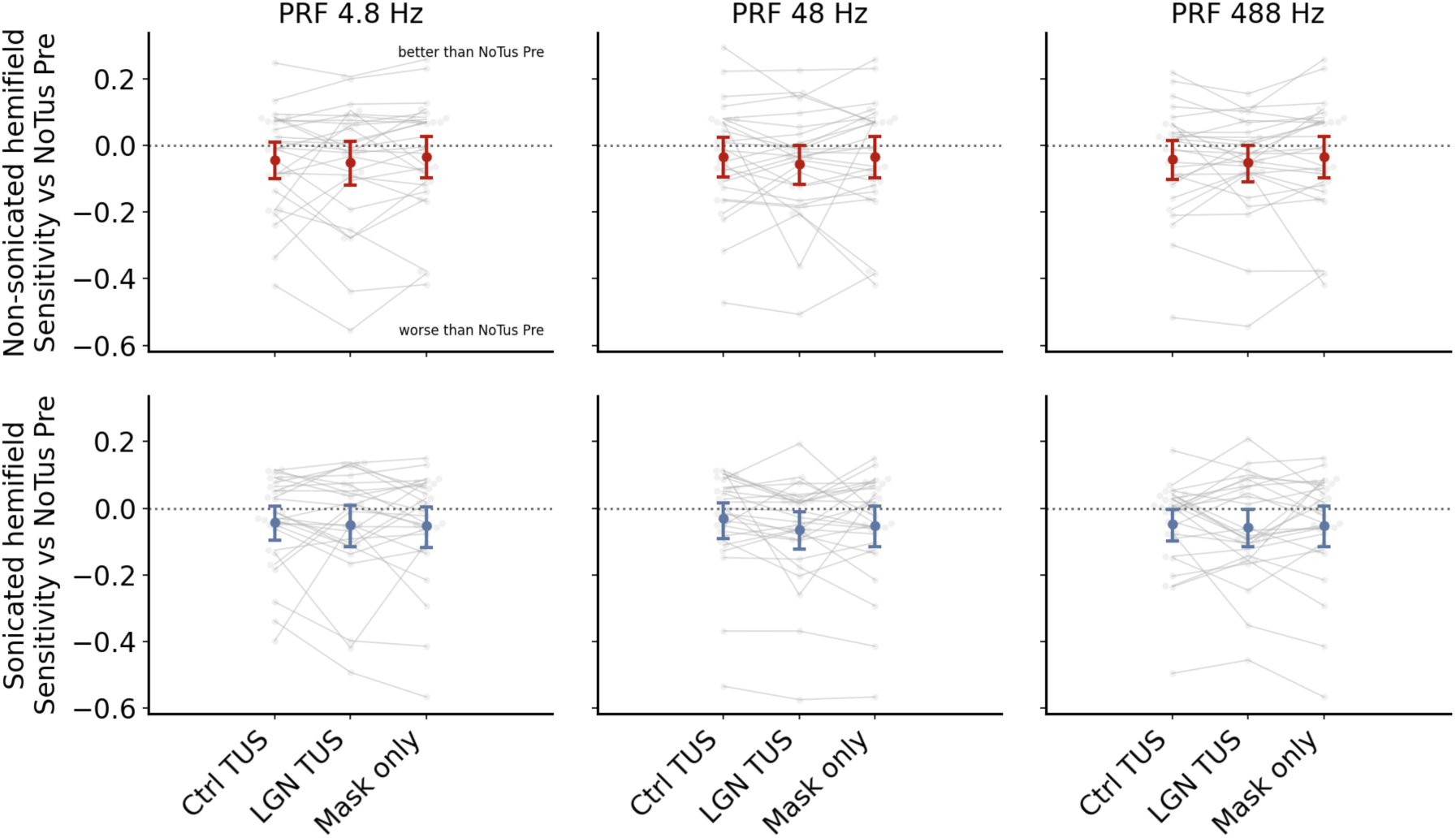
Non-significant effect of LGN TUS on visual perceptual sensitivity. Behavioral thresholds normalized to pre-TUS baseline for the control (Ctrl) TUS, LGN-TUS, and mask only conditions, separately for each PRF (columns) and hemifield (rows; active stimulation on bottom). Larger colored markers show means and 95% confidence intervals, with smaller grey markers depicting individual subject values.

We examined whether the variance in the interaction between TUS target and hemifield could be associated with the variability in TUS dose (summarized as the mean simulated I_SPPA_ in the LGN). The fits of the linear model for each PRF are shown in **Figure 5**. At 4.8 Hz, higher mean I_SPPA_ in the LGN was associated with the interaction value (slope = -0.0188, r = -0.412, R² = 0.169, p = 0.037). However, this relationship passes through zero, with a lower dose predicting a larger LGN-Ctrl TUS difference in the sonicated hemifield, and a higher dose predicting a larger LGN-Ctrl TUS difference in the Non-sonicated hemifield. There was no clear evidence of a dose relationship at the higher PRFs. At 48 Hz, the association was near zero (slope = 0.00347, r = 0.078, R² = 0.006, p = 0.704), and at 488 Hz the association was also weak (slope = -0.00591, r = -0.094, R² = 0.009, p = 0.648).

**Figure 5:**
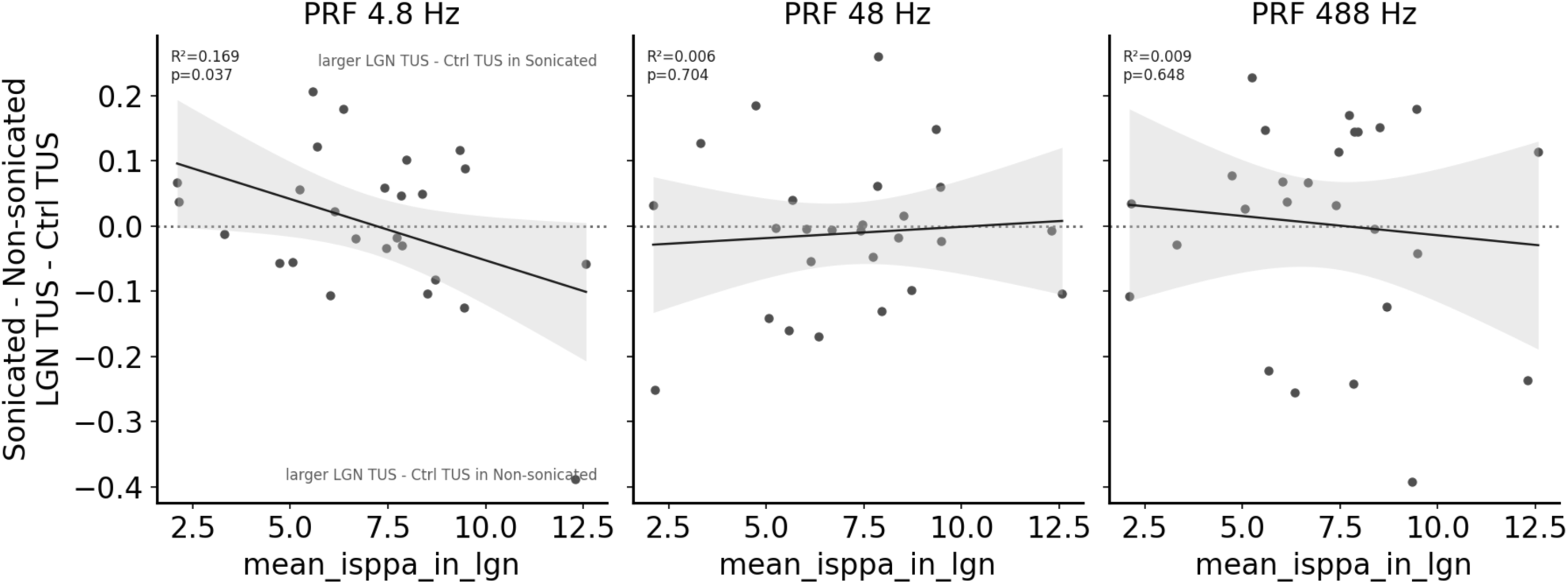
Dosimetry – behavior. Mean simulated I_SSPA_ at the LGN vs the hemifield x depth interaction effect. The interaction was plotted per individual and fit with a linear regression (black line, grey line shows 95% confidence interval) for each PRF. P-value and R^2^ for each fit shown as inset.

### Effect of LGN TUS on concurrent SSVEP amplitudes

We then tested the hypothesis that LGN TUS would disrupt or enhance EEG steady-state visual-evoked potential responses in the contralateral (sonicated) hemifield, and that this effect may be PRF-specific. The 2F SSVEP amplitudes normalized to the pre-TUS baseline are shown in **Figure 6** for each of the online experimental conditions. Notably, for all conditions displayed in **Figure 6**, all 95% CIs overlap with each other and the unity line, which indicates that there is little evidence for consistent differences between any active or control TUS conditions. A repeated-measures ANOVA revealed no evidence of a main effect of PRF (WTS(2) = 0.13, p = 0.937), TUS target (WTS(2) = 2.499, p = 0.126), hemifield (WTS(1) = 3.577, p = 0.070), or the PRF × target interaction (WTS(2) = 1.491, p = 0.488). No interactions involving hemifield were significant: PRF × hemifield (WTS(2) = 0.298, p = 0.861); target × hemifield (WTS(1) = 0.091, p = 0.763), nor the 3-way PRF × target × hemifield interaction (WTS(2) = 0.160, p = 0.923). Overall, the results of this analysis do not support the presence of target-specific, hemifield-specific, or PRF-specific effects of LGN-TUS on the amplitude of the SSVEP.

**Figure 6:**
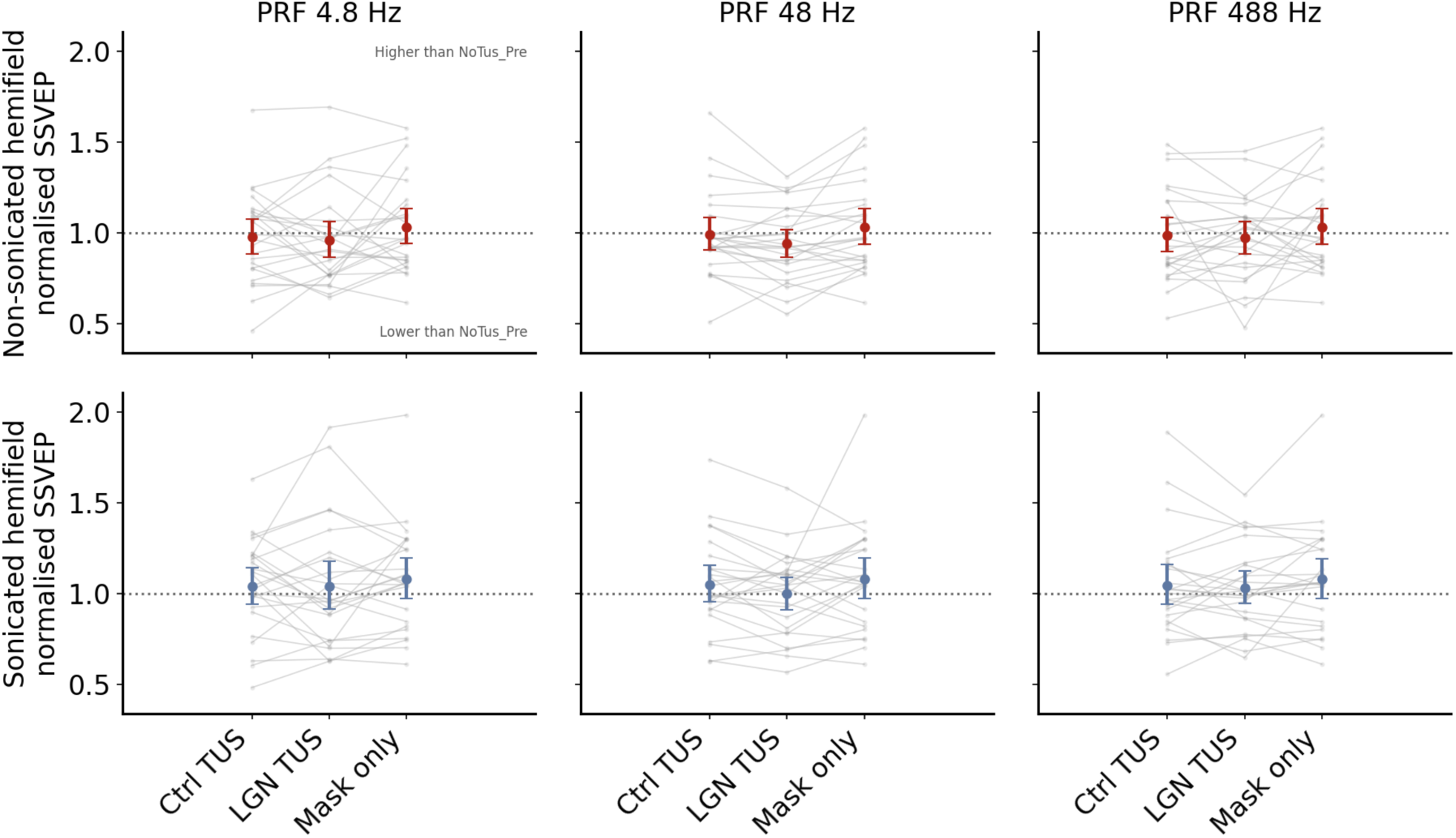
Non-significant online effect of LGN TUS on EEG visual-evoked response amplitude. 2F SSVEP amplitude in µV, normalized as fraction of pre-TUS, for the control (Ctrl) TUS, LGN TUS, and mask only conditions, separately for each PRF (columns) and hemifield (rows; active stimulation on bottom). All formatting is the same as Figure 3.

**Figure 7** shows the interaction between hemifield and TUS target against the simulated mean I_SPPA_ within each participant’s LGN ROI for the SSVEP amplitude. There was no evidence that mean I_SPPA_ in LGN was associated with the interaction term for any PRF. At 4.8 Hz, the fitted association was negative but small (slope = -0.0219, R^2 = 0.063, p = 0.357). At 48 Hz, the fitted association was positive but similarly weak (slope = 0.0162, R^2 = 0.065, p = 0.357). At 488 Hz, the association was effectively absent (slope = -0.00285, R^2 = 0.001, p = 0.878). Overall, there was little evidence of a systematic shift in the sonicated-versus-non-sonicated LGN TUS effect as a function of TUS dose.

**Figure 7:**
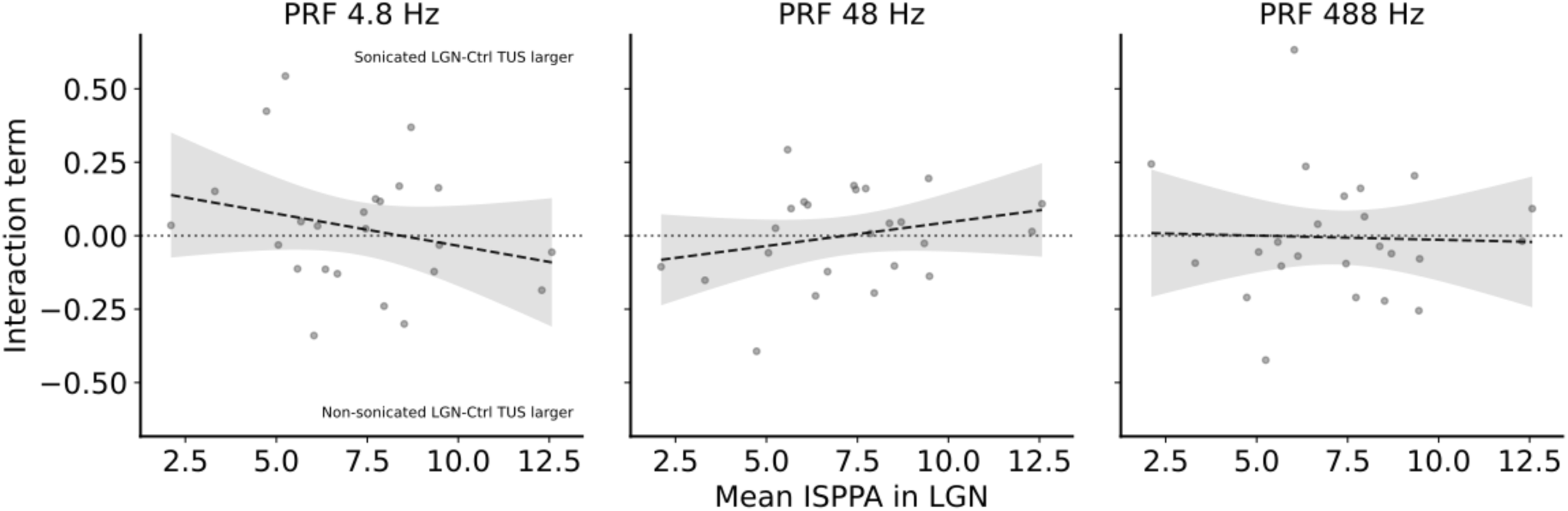
Minimal dose-response relationship between TUS and SSVEP amplitude. Dosimetry for the SSVEP amplitude hemifield x TUS target interaction term. The OLS linear fit is shown as a dashed line, with the 95% CI on the model fit shown shaded in grey.

### Effect of LGN-TUS on SSVEP response latency (phase)

We next tested the hypothesis that TUS could modulate the latency of the visual-evoked response, which would be apparent as a change in phase of the SSVEP response harmonic. The phases of the 2F responses are shown in **Figure 8**. Here, there is little evidence for a consistent online effect of TUS on the response phase, save for a trend towards a slower response (relative to the pre-TUS baseline) in the LGN-TUS condition for the non-sonicated hemifield at a PRF of 487.5Hz (note again that a veridical TUS effect would be more apparent in the contralateral sonicated hemifield). In addition, a repeated-measures ANOVA revealed a marginally significant interaction between TUS target and hemifield, providing weak evidence that the target affected response phase differently between the two hemifields (WTS (1)= 4.374, p = 0.047). There was no main effect of PRF (WTS(2) = 0.417, p = 0.812), TUS target (WTS(1) = 1.190, p = 0.275), or hemifield (WTS(1) = 0.036, p = 0.849). There was also no evidence for a PRF × hemifield interaction (WTS(2) = 2.391, p = 0.303) or a 3-way PRF × target × hemifield interaction (WTS(2) = 0.690, p = 0.708). The PRF × stimulation interaction was also non-significant (WTS(2) = 5.158, p = 0.106).

**Figure 8:**
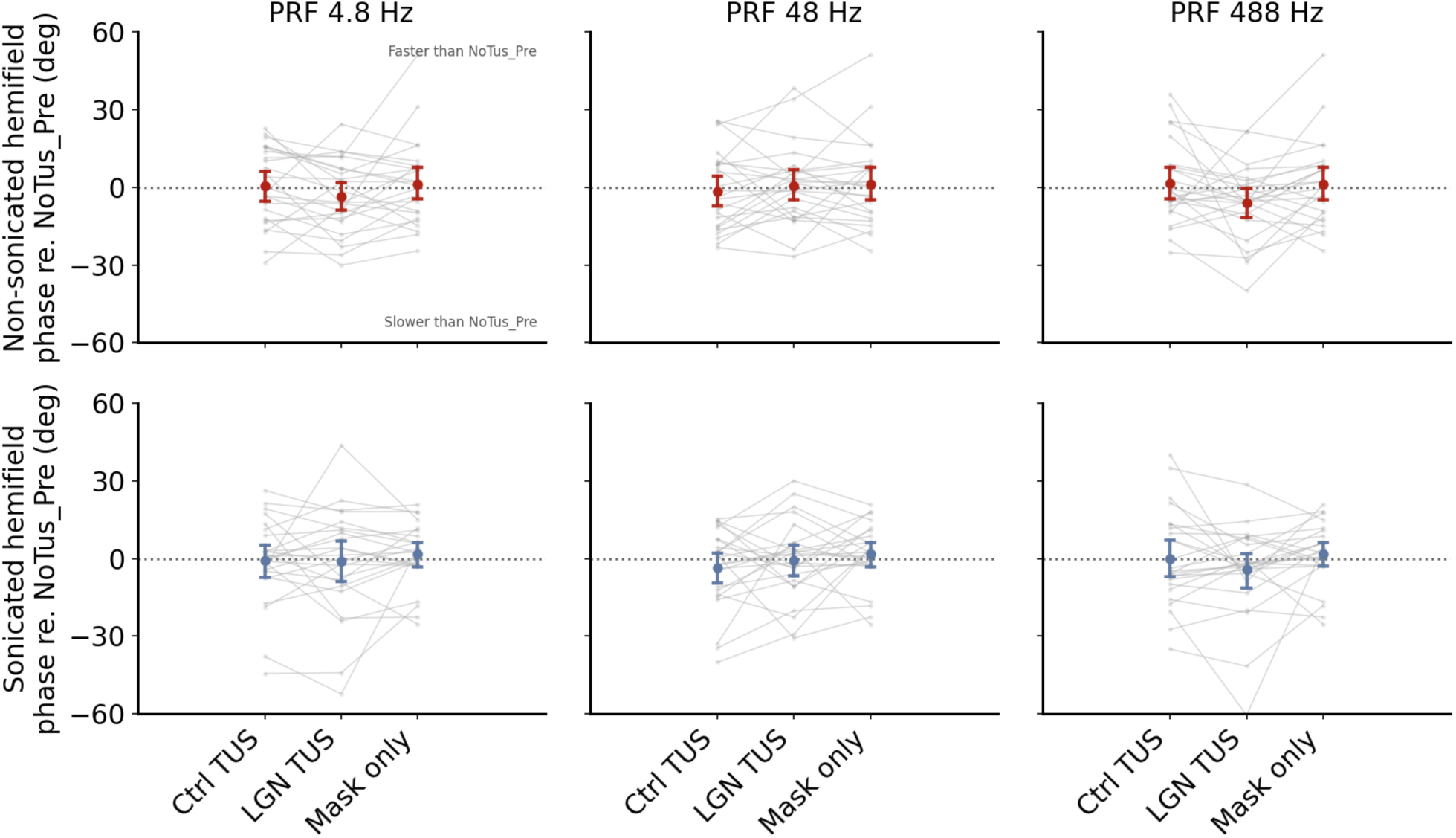
Non-significant online effect of LGN TUS on EEG visual evoked response latency. 2F SSVEP phase in degrees, normalized by subtraction from pre-TUS, for the control (Ctrl) TUS, LGN TUS, and mask only conditions, separately for each PRF and hemifield of origin. All formatting is the same as Figure 6.

**Figure 9** shows the dosimetry effect for phase. There was no evidence of a dose-response relationship at any PRF tested. At 4.8 Hz, the fitted association was weak and negative (slope = -0.85, R^2 = 0.041, p = 0.542); at 48 Hz, it was weak and positive (slope = 0.84, R^2 = 0.015, p = 0.572); and at 488 Hz, it was again weak and negative (slope = -1.00, R^2 = 0.038, p = 0.542). Overall, this figure indicates that variation in mean I_SPPA_ in LGN did not reliably explain between-participant variation in the 2F phase interaction effect.

**Figure 9:**
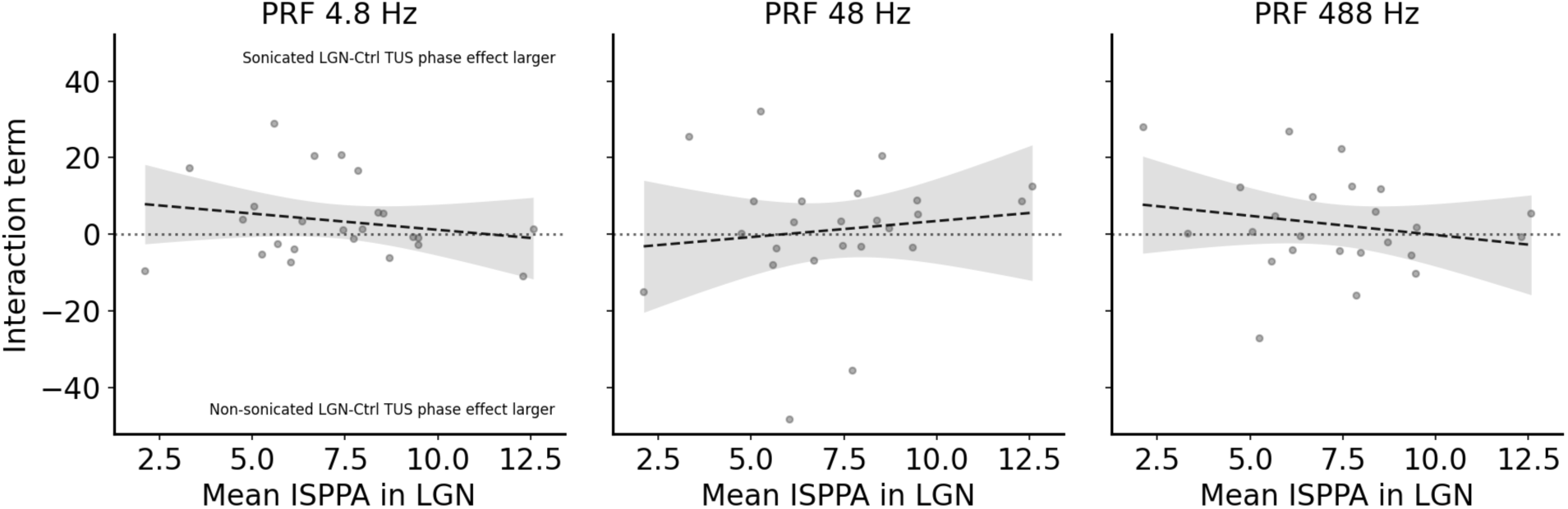
No dose-response relationship between TUS and SSVEP response latency. Dosimetry for the SSVEP phase hemifield x TUS target interaction term. The OLS linear fit is shown as a dashed line, with the 95% CI on the model fit shown shaded in grey.

### No evidence of a cumulative offline effect

Although we failed to detect any online effect of TUS, we reasoned that the effects of TUS across the entire experiment could accumulate and produce changes in the post-TUS condition. These would be expected to be observable in the sonicated (contralateral) hemifield. **Figure 10A** shows the sensitivity thresholds for the pre-TUS time point, interleaved no-TUS time-point, and the post-TUS time-point, normalized to the pre-TUS condition. Note that in all of these time-points only the auditory mask was played during the trial. In both hemifields, there was a clear trend towards sensitivity being worse in the conditions following the pre-TUS condition. A repeated-measures ANOVA showed a significant main effect of time-point (WTS(2) = 9.67, p = 0.0223), indicating that normalized sensitivities decreased across the Pre, Control, and Post time-points. However, no main effect was observed for hemifield (WTS(1) = 0.12, p = 0.7245), and the hemifield by time-point interaction was not significant (WTS(2) = 6.31, p = 0.0654), indicating a non-specific reduction in performance more consistent with participant fatigue than a hemifield-specific effect of TUS neuromodulation. As a dosimetry test (**Fig. 10B**), we tested whether the simulated mean I_SPPA_ in the LGN was associated with the pre-post and sonicated - Non-sonicated interaction term. There was no evidence of a linear association between dose and this interaction (slope = -0.00538, r = -0.121, R² = 0.015, p = 0.557).

**Figure 10:**
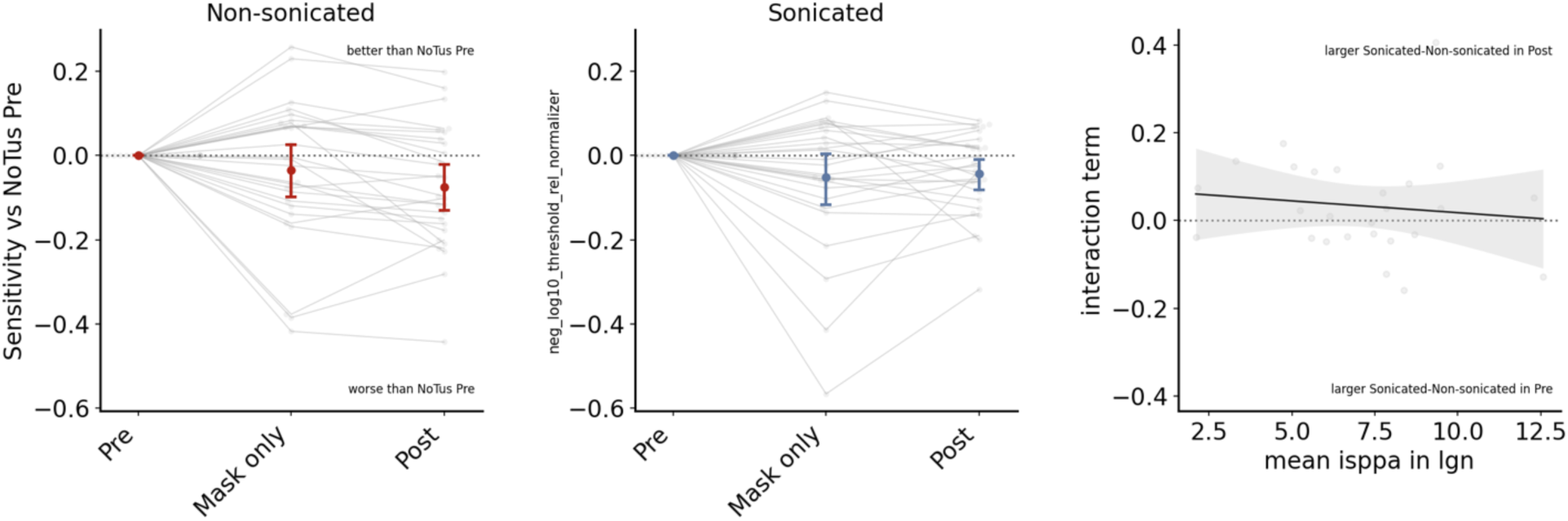
Non-significant offline effects of LGN TUS on perceptual behavior. **A.** Normalized behavioral sensitivity for the conditions where no TUS was actively delivered (pre-TUS, mask only, and Post-TUS), separately for the sonicated and non-sonicated hemifields. **B**. Scatter plot and linear regression of estimated TUS intensity at LGN vs. Difference in performance between non-sonicated and sonicated hemifields at post-TUS timepoint.

**Figure 11A** shows the SSVEP amplitude for the no TUS conditions as normalized to the Pre-TUS baseline condition. As all confidence intervals contain the reference value, it would appear there is little evidence of an offline effect of TUS. Indeed, a repeated-measures ANOVA reported no significant main effect of hemifield (WTS(1) = 1.779, p = 0.194), or time-point (WTS(2) = 0.945, p = 0.653), and no hemifield × time-point interaction (WTS(2) = 0.905, p = 0.664). There was no dose-response relationship between SSVEP amplitude and offline LGN TUS (**Fig. 11B**).

**Figure 11:**
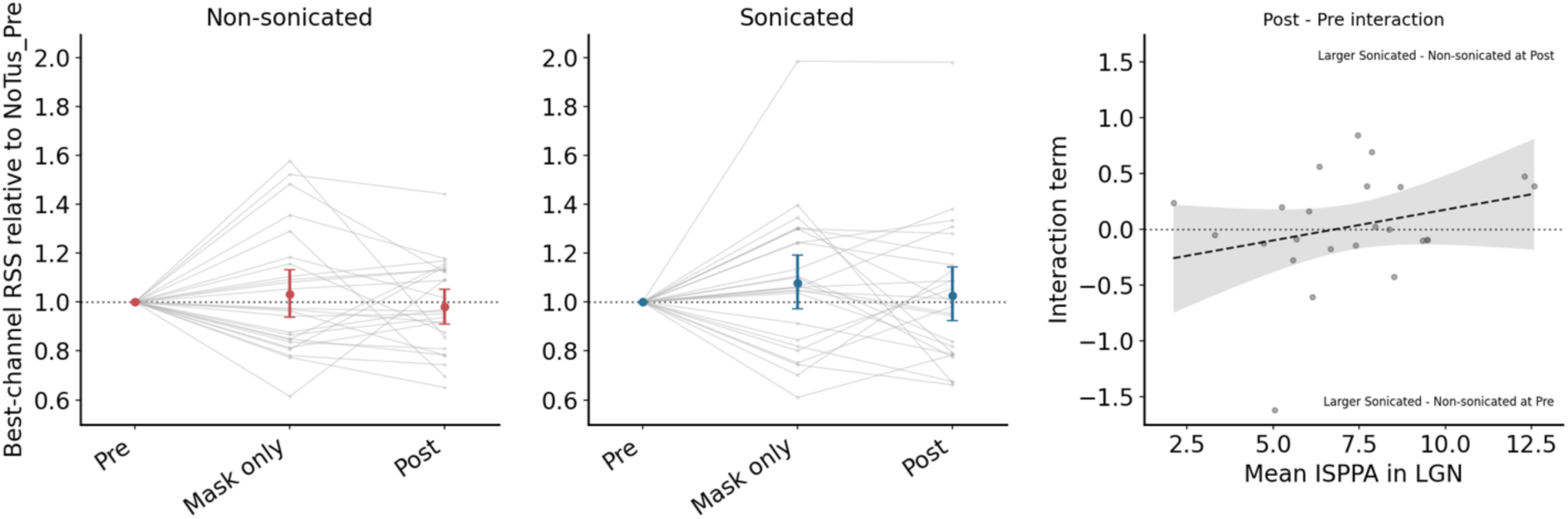
Non-significant offline effect of TUS on SSVEP response amplitude. **A.** Normalized SSVEP amplitude for the conditions where no TUS was actively delivered (pre-TUS, mask only, and Post TUS), separately for the sonicated and non-sonicated hemifields. **B**. Scatter plot and linear regression of estimated TUS intensity at LGN vs. Difference in the change in SSVEP amplitude Pre to Post between non-sonicated and sonicated hemifield.

For the SSVEP response latency, shown in **Figure 12A**, there was again no clear evidence for an offline effect on the basis of the confidence intervals for each condition. In a repeated-measures ANOVA on the 2F phase, there was no significant main effect of hemifield (WTS(1) = 0.424, p = 0.520), no significant main effect of time-point (WTS(2) = 0.026, p = 0.988), and no hemifield × time-point interaction (WTS(2) = 0.001, p = 1.000). Additionally, there was no evidence that the mean I_SPPA_ in LGN was associated with the pre-post and sonicated – non sonicated interaction term (**Fig. 12B**). The fit was effectively flat (slope = 0.264, intercept = -1.05), with a negligible association strength (r = 0.034, R^2^ = 0.0012) and was not significant (p = 0.875).

**Figure 12:**
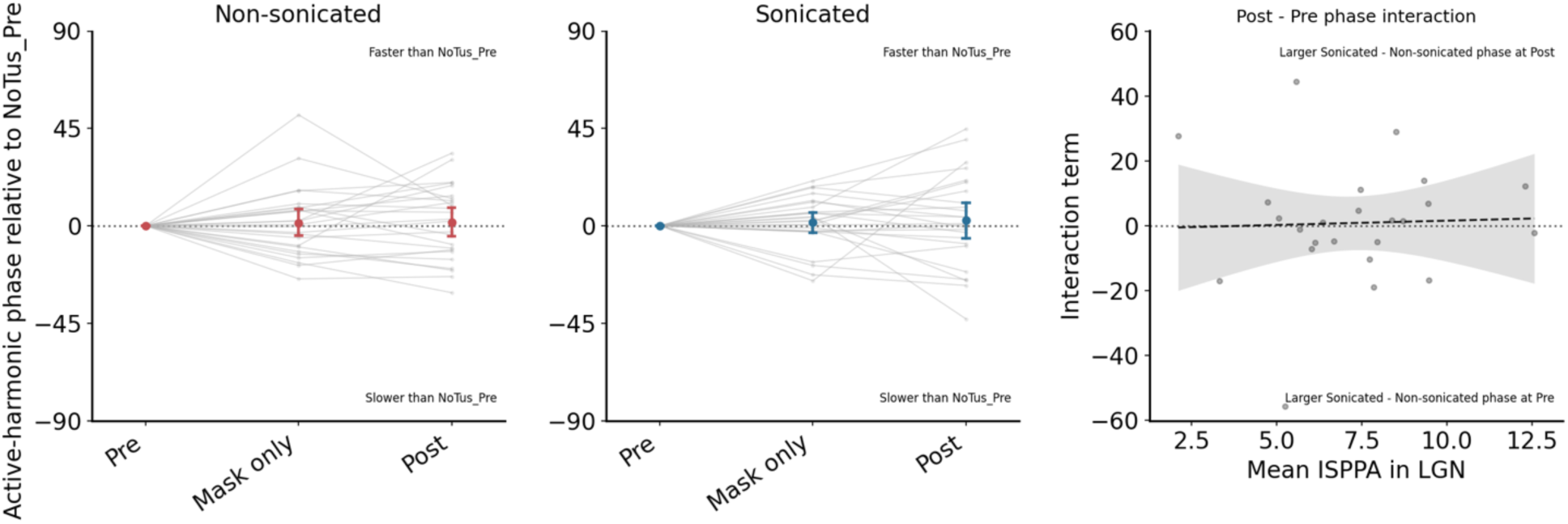
Minimal offline effect of TUS on SSVEP response latency. **A.** Normalized 2F response phase for the conditions where no TUS was actively delivered (pre-TUS, mask only, and Post TUS), separately for the sonicated and non-sonicated hemifields. **B**. Scatter plot and linear regression of estimated TUS intensity at LGN vs. Difference in the change in SSVEP phase Pre to Post between non-sonicated and sonicated hemifield.

### Assessment of targeting accuracy and statistical power

The 3D position of the TUS transducer was tracked online in Brainsight, allowing us to more accurately simulate where the TUS beam landed in individual participants. We plotted individual participant LGN spheroid positions on top of the across-participant average TUS beam volume in **Fig. 13**, with each panel showing the two long-axis and the short-axis profiles of the beam.

**Figure 13:**
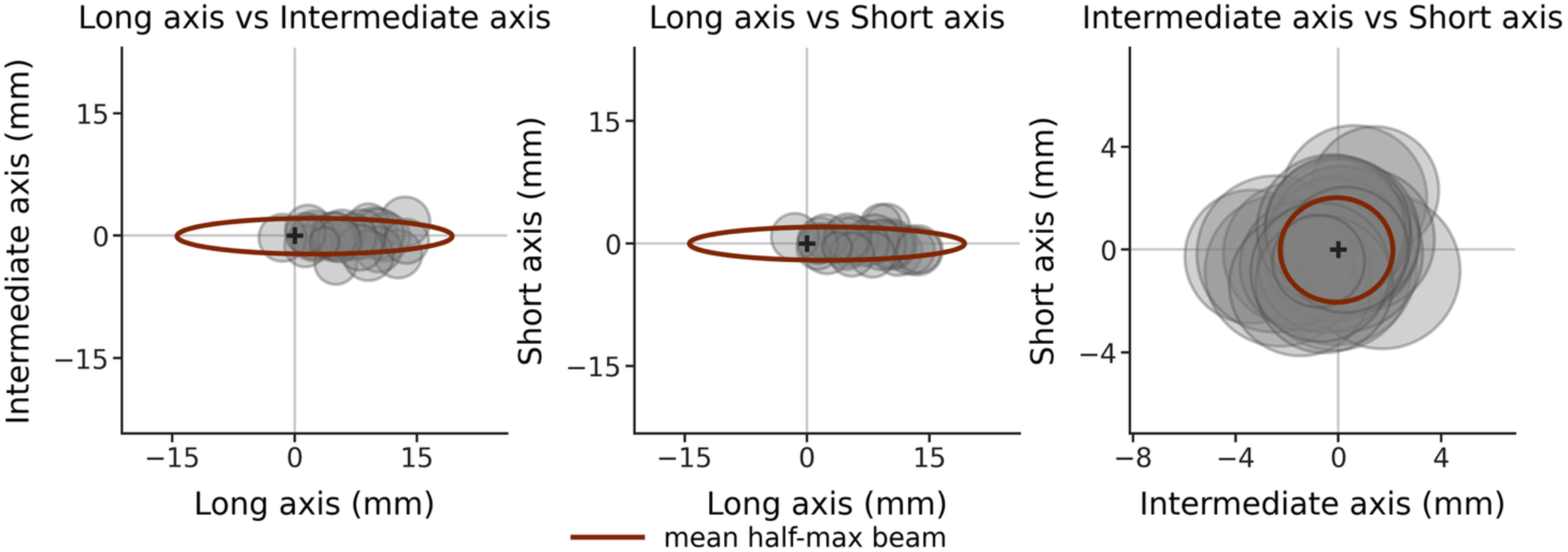
Demonstration of TUS targeting accuracy across participants. The spheroid LGN volumes for each participant are plotted as grey circles on the simulated FWHM TUS beam (red dotted oval/circle). Left, middle: long-axis profiles; right: short axis profile. The black cross depicts the peak of the TUS focus. The FWHM TUS volume overlapped with LGN in all but two participants. The 70 mm steering depth was somewhat shallow, indicating that the peak of the TUS focus was slightly shallow compared to the LGN

A sensitivity analysis conducted in G*Power indicated that, with N=25, α=.05, and 80% power, the study could detect a within-subject interaction effect of f≥0.264. We assumed a correlation of r=.591 between repeated measures, which was calculated based on the correlation between the mean of the hemifield x TUS-depth interaction terms for each PRF level. Thus, for SSVEP amplitude effects, the study was adequately sensitive to detect approximately medium or larger interaction effects.

## Discussion

This study aimed to interrogate the role of the PRF in TUS neuromodulation effects, using the early visual system, the LGN, as a test-bed, and simultaneous recordings of SSVEPs and contrast sensitivity as a readout. Our primary hypotheses were that TUS to the LGN would 1) have larger impacts on SSVEP responses and perceptual behavior than TUS to a non-visual control site; 2) have larger impacts on the hemifield contralateral to stimulation (the sonicated hemifield) compared to the ipsilateral non-sonicated hemifield, and 3) have differing effects depending on PRF. In contrast to several prior experiments from our lab and others (Fry, Ades, and Fry 1958; Mohammadjavadi et al. 2022; Webb et al. 2023, 2022; Martin et al. 2025), including a pilot experiment from our research group (R. Ash et al. 2025), our experiment failed to find evidence for any of the three hypotheses. The EEG and perceptual behavior measures were, on average, stable across each of the experimental conditions. A dosimetry analysis based on simulations from the measured transducer position during TUS showed that LGN was within the FWHM of the focus in the majority of participants and failed to detect any stronger effect in the participants with higher-intensity LGN stimulation.

Several strengths of our study lend credence to the findings. Steady state VEP frequency tagging (Norcia et al. 2015) allowed for a simultaneous control stimulus in the contralateral hemisphere, allowing estimates of the specificity of TUS effects. Our measures had high signal-to-noise ratio, and have previously been shown to have exceptional across-time stability (R. T. Ash, Nix, and Norcia 2024). We also employed a psychophysical task to capture any subtle changes in perceptual sensitivity. The control-depth TUS condition accounts for any residual non-specific TUS effects, and a white-noise mask rendered the auditory co-stimulation of TUS near-imperceptible (**Supp. Figure 2**). Together, these factors protect our inferences from auditory and expectancy confounds that have complicated the interpretability of some previous TUS experiments (Guo et al. 2018; Kop et al. 2024; Sato, Shapiro, and Tsao 2018; Fong et al. 2025). Although we do not have ground-truth proof of accurate targeting, measurements of our group and others have shown that simulation accurately predicts the location of the focal spot (Martin, Jaros, and Treeby 2020). Despite all of these experimental optimizations, we did not identify consistent neuromodulatory effects of TUS.

### Explanations for the absence of a TUS effect

It is unclear why we failed to detect TUS neuromodulation effects. In contrast to dozens of studies showing neuromodulation effects ex vivo in animals and humans across many brain areas (Blackmore et al. 2019; Pellow, Pichardo, and Pike 2024; Sarica et al. 2022), including the LGN (Martin et al. 2025; Webb et al. 2023; Mohammadjavadi et al. 2022). We address three potential reasons for the lack of robust neuromodulation effects in turn: 1) Insufficient mechanical or thermal dose; 2) inaccurate targeting; 3) online effects vs. offline effects.

It is possible that the absence of TUS neuromodulation effects reported is due to an inadequate mechanical or thermal dose. We cannot exclude the possibility that a higher mechanical dose would have produced stronger neuromodulation effects, although we used intensities (68 W/cm^2^ free-water, estimated 8 W/cm^2^ on average at the target) at the high range of existing human studies, and similar to many animal studies (Darmani et al. 2022). We note here that BabelBrain simulations estimate higher *in situ* intensities compared to K-Plan, the other commonly used TUS simulation software (Brandts et al. 2026). Another compelling possibility is that the thermal dose we achieved was insufficient to acutely modulate thalamic activity. The thalamus is known to be highly sensitive to even small 0.5-1 ℃ rises in temperature (Darrow et al. 2019) as are some other areas (Owen, Liu, and Kreitzer 2019). Fry et al (1958) and Mohammadjavadi (2022) used higher intensities and higher duty cycles than what are commonly used in human studies, so it is likely that the temperature increased at the target in these studies.

It is possible that the absence of TUS effects in our data is also due to inconsistent targeting of the LGN. The LGN is a small ∼ 0.5 cm diameter structure that is deep in the brain, ∼6 cm from the surface. In most participants, the ear is lateral to the LGN, such that the transducer must be positioned at an angle above the ear to target the LGN. This angle increases the transducer-to-LGN distance and the incident angle on the skull, which increases reflection and acoustic lensing. The TUS focus with a 4 element 500 kHz transducer is approximately 0.5 x 0.5 x 2 cm FWHM. Therefore, even a small error in positioning or angle will lead to missing the target. Infrared neuronavigation like the Brainsight system we use here is known to have ∼0.5 cm of error (Phipps et al. 2024). Thus, it is possible that we missed the LGN in a subset of participants due to scalp-surface neuronavigation registration errors, which are difficult to assess directly with our current equipment. Furthermore, previous hydrophone measurements of TUS through the skull have confirmed that simulations estimate the position of the focus well, but the dose less well (Martin, Jaros, and Treeby 2020), which may contribute to errors in the dose estimates used in our dosimetry analysis. Indeed, with a ground-truth dose and target engagement assessments like MR-ARFI, an effect may have been revealed in our data.

A third possibility for the lack of TUS effects is that ‘offline’ effects are more robust than ‘online’ effects. Online effects refer to the immediate neuromodulation that occurs coincident with TUS, which our experiment was primarily designed to investigate. Offline effects, in contrast, refer to the delayed effects that emerge minutes and hours after stimulation (Bault, Yaakub, and Fouragnan 2024). Offline effects have been repeatedly shown in primates and humans with resting state functional MRI (Fouragnan et al. 2019; Folloni et al. 2019; Verhagen et al. 2019; Yaakub et al. 2023; Atkinson-Clement, Alkhawashki, et al. 2024) and other methods (Webb et al. 2023) and can last from minutes up to hours after stimulation. Our experiment was not optimized to detect offline effects, but the Pre/Post blocks of trials would have been expected to detect any gross cumulative effect of the ∼30 minutes of ultrasound that participants received during the TUS blocks. No hemifield-specific change in perceptual sensitivity or SSVEP responses was observed over the course of the experiment, but we acknowledge that effects may have manifested after the session was concluded, that we would not have captured.

### Relation to previous TUS LGN findings

Our experiment failed to replicate the perceptual effects of ultrasound to LGN reported in non-human primates (Webb et al. 2023). In these studies, TUS to the LGN in awake behaving monkeys (targeting confirmed with MR thermometry) caused a parameter-dependent enhancement or suppression of visual perceptual sensitivity that was lateralized as expected given the anatomy of the visual pathway. These effects were observable for an enduring period following stimulation. Compare this to the current study, in which even with similar TUS sonication parameters, we did not observe robust online or short-term offline effects of TUS on behavior. In addition to targeting, perhaps these effects were also due to a combination of thermal neuromodulation and/or auditory co-stimulation, but the true reasons for this disparity await further experimental clarification.

Our results also disagree somewhat with a recent well-controlled study showing that TUS to the LGN can either suppress or enhance visual-evoked functional MRI BOLD responses in visual cortex, using sonication parameters similar to the TUS parameters of some of the conditions we use here and an active control site (Martin et al. 2025). Three potential explanations for this dissociation include 1) robust neuromodulation effects were only observed in a small subregion of visual cortex in their study – since SSVEPs measure synchronized activity from large swaths of visual cortex, a change in activity in a small subregion may be undetectable in the EEG due to the unchanged activity in the rest of visual cortex. 2) Though we used a similar PRF in one of our conditions (4.8 Hz vs 5 Hz) their study used a much longer pulse length (300ms). Their study had a smaller TUS focus that completely overlapped with the LGN due to a high number of elements (256 vs 4 here) while our focus overlapped with LGN and several nearby structures, particularly the optic radiations and thalamic reticular nucleus, the stimulation of which could interact with stimulation effects at the LGN; and 3) their in-scanner phase array system allowed much more precise targeting, so they may have been more consistently delivering TUS to LGN vs our neuro- navigated study

An important less-recognized implication of the failure to replicate the pilot experiment’s results in the current experiment is that the auditory confound can generate spuriously lateralized differences between conditions. In general, a lateralized difference would be strong evidence for a specific effect in this paradigm, due to the lateralized hemispheric anatomy of the brain. However, in the current experiment the position of the transducer on the left side of the head and the lateralized sounds generated by it could bias attention to the visual stimuli. The auditory confound can influence response through attentional effects: drawing attention to the sonicated side (Hidaka and Ide 2015), and placebo- expectancy effects (receiving a ‘treatment’ on the left side (Sterzer et al., 2008; Schienle et al., 2014; Tiraboschi et al., 2019)). Auditory co-stimulation can be minimized with auditory masks (Braun et al. 2020; Johnstone et al. 2021; Liang et al. 2023) and ramped waveforms when possible (Mohammadjavadi et al. 2019). In the near future, phase array transducers will allow a more robust control TUS condition that is defocused (Caulfield et al. 2025) or targets an inactive site (e.g. the lateral ventricle (Atkinson-Clement, Kaiser, et al. 2024)) which precisely matches the sensory co-stimulation of active stimulation without focally depositing ultrasound in the brain, avoiding potentially confounding CNS effects.

### Limitations and future priorities

We want to note additional limitations of the study that can be improved on in the future. First, although we advised participants to fixate at a central cross, and participants performed a task that required equal attention to both hemifields, we did not track eye movements, so eye movements in some participants could have increased the variability of our results. Another limitation is that the optimal LGN trajectory would require the transducer path to go through the ear, which is not feasible to pass through given the presence of air pockets. With our 4-element annular transducer, we could not beam-steer laterally, so instead we had to place the transducer at an angle on the head (5-10°). This decreased the amount of TUS that made it to the target due to reflections and refractions by the skull. Future experiments with phase arrays will allow beam steering to targets like the LGN without the need for angled transducer positions and allow for phase aberration correction. A related limitation is that our active control condition may have inadvertently stimulated visual-related pathways, either through modulation of the optic radiations or other noncanonical pathways. This active control site was limited by what we could achieve with our 4-element transducer. In contrast, phase array transducers allow control TUS site targeting to the lateral ventricle (Atkinson-Clement et al., 2024) or a defocused control that matches the auditory effect but does not deliver a focus of ultrasound pressure in the brain (Caulfield et al., 2025). Although note that the lack of differences between the active and no-TUS conditions makes this a non-issue for the current results.

Another limitation is that our CTX500 transducer had a maximum depth of 70 mm, and in some participants due to head size and trajectory angle, the TUS focus was not able to reach deep enough to hit the LGN. As the TUS beam is cigar-shaped and extended in the axial dimension, our simulations show that the LGN was in the >70% of max intensity focus for all participants. Nevertheless, future experiments should use a transducer with a larger maximal depth, adjust the depth of sonication based on individual simulations, and ideally use ARFI measurements to quantify dose variability. Another limitation is that we used the same free-water TUS intensity in all participants, despite the well-known variability in skull attenuation across individuals. Some participants received significantly higher TUS doses at the target than others (range 8.5-20 W/cm^2^ I_SPPA_ as per BabelBrain). We made this experimental choice to balance maximizing TUS intensity and safety. If we had decided to match all participants at a high intensity, the subset of participants with high attenuation would require high powers and may have had significant heating in the skull or scalp. If we had decided to match participants at a low intensity, we may not meet the TUS dose needed to have significant online effects.

Another potential source of variability in the experiments is the neuronavigation. We made significant effort to optimize the fidelity of our fiducial placement and during the validation step we had a conservative threshold of 2 mm error on the sides, front, and top of the head. We also monitored the TUS focus position closely during experiments to keep it within 0.5 mm of the target. Given that the size of the focus is 5 mm in the lateral direction, if our error is < 4 mm, then we should be at least partially modulating LGN in the reported experiments, which was supported by our post-hoc simulations and dosimetry **(Fig. 13)**. Additional potential sources of error in neuronavigation include calibration of the TUS transducer and errors in MRI segmentation. There are no standards currently to our knowledge about optimal neuronavigation in TUS experiments, and this is an essential place for experimental improvement. Newer technologies with machine vision are expected to provide an order of magnitude improvement in neuronavigational precision (Chiurillo et al. 2023). When experiments can be performed in-scanner, MR-ARFI, which visualizes the TUS focus in situ, will provide a new gold-standard method to verify the TUS focality and targeting accuracy and estimate dose (Mohammadjavadi et al. 2025).

## Conclusion

Our results provide a cautionary note on the robustness and effect sizes of neuro-navigated TUS in studies of the human visual system. It will be essential in the future to confirm the inefficacy of LGN TUS with proof-positive ARFI-confirmed targeting, to see if similarly rigorous paradigms fail to show effects in other sensory systems, and to develop rigorous measures of TUS effects that are generalizable across brain areas.

