## Supplementary material for "Effects of transcranial focused ultrasound stimulation to human lateral geniculate nucleus on visual perception and steady-state visual evoked potentials"

### Supplement 1: Hydrophone ultrasound characterization

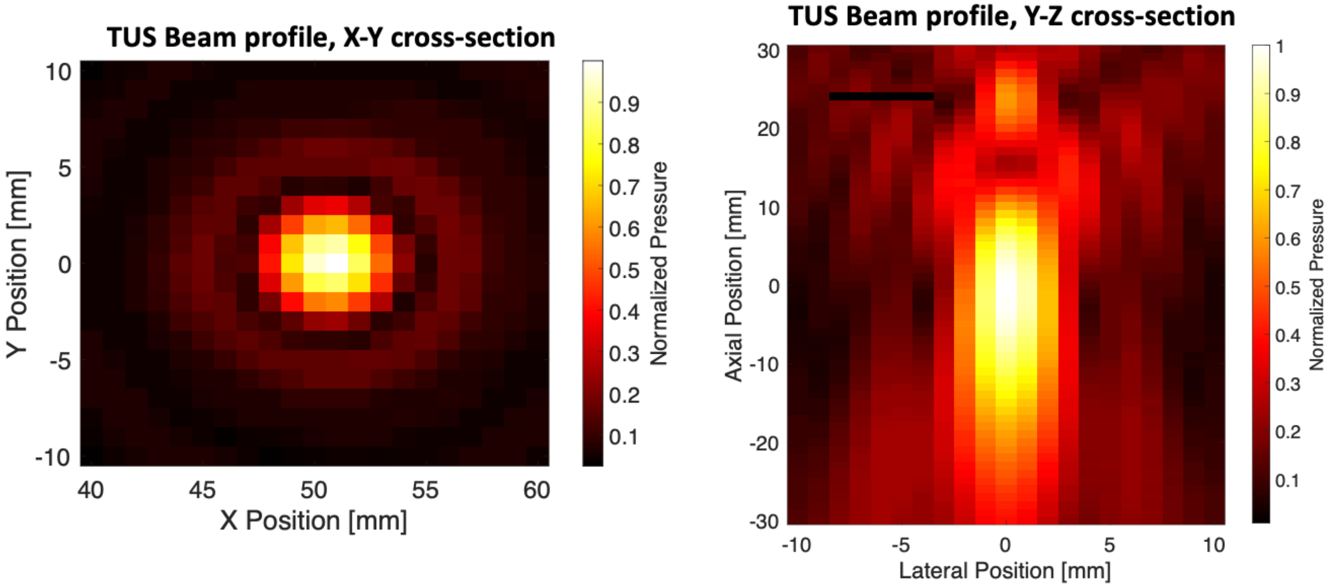


**Supplementary Figure. 1** Hydrophone measures of our NeuroFUS CTX-500 transducer beam volume. Measured at 10W/cm$^2$ $I_{SPPA}$ output, steered to 60mm depth. **Left panel**: X-Y view shows lateral cross-section of TUS beam profile. **Right Panel**: Y-Z view shows axial cross-section of beam profile.

**Supplement 2: Auditory masking**

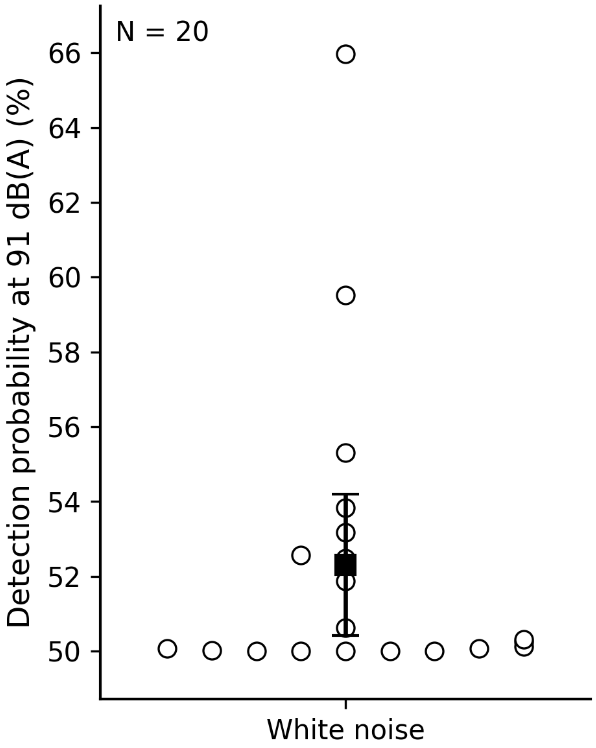


**Supplementary Figure 2:** Predicted detection probability of LGN-TUS at under a 91 dB(A) white-noise masker. Open circles show individual participants. The black square shows the group mean, and error bars show the 95% confidence interval. Detection probability is plotted as percent correct from a fitted psychometric function from a supplementary two-interval forced-choice experiment that 20 participants returned for. In this experiment, the loudness of the mask was varied to determine the full shape of the LGN-TUS detectability with a 68 W/cm$^2$ $I_{SPPA}$ and a similar transducer placement above the ear.
